# Creld2 function during unfolded protein response is essential for liver metabolism homeostasis

**DOI:** 10.1101/2020.01.28.923136

**Authors:** Paul Kern, Nora R. Balzer, Franziska Bender, Alex Frolov, Klaus Wunderling, Jan-Peter Sowa, Lorenzo Bonaguro, Thomas Ulas, Christoph Thiele, Joachim L. Schultze, Ali Canbay, Reinhard Bauer, Elvira Mass

**Affiliations:** Developmental Biology of the Immune System, Life & Medical Sciences (LIMES) Institute, University of Bonn, 53115 Bonn, Germany; Developmental Genetics & Molecular Physiology, Life & Medical Sciences (LIMES) Institute, University of Bonn, 53115 Bonn, Germany; Biochemistry & Cell Biology of Lipids, Life & Medical Sciences (LIMES) Institute, University of Bonn, 53115 Bonn, Germany; Department of Gastroenterology, Hepatology and Infectious Diseases, University Hospital Magdeburg, 39120 Magdeburg, Germany; Department of Medicine, Ruhr University Bochum, University Hospital, Knappschaftskrankenhaus Bochum, 44892 Bochum, Germany; Genomics and Immunoregulation, Life & Medical Sciences (LIMES) Institute, University of Bonn, 53115 Bonn, Germany; Platform for Single Cell Genomics and Epigenomics at the German Center for Neurodegenerative Diseases and the University of Bonn, 53127 Bonn, Germany

**Keywords:** ER stress, UPR, NASH, liver steatosis

## Abstract

The unfolded protein response (UPR) is associated with the hepatic metabolic function, yet it is not well understood how endoplasmic reticulum (ER) disturbance might influence metabolic homeostasis. Here, we describe the physiological function of Cysteine-rich with EGF-like domains 2 (Creld2), previously characterized as a downstream target of the ER-stress signal transducer Atf6. To this end we generated *Creld2*-deficient mice and induced UPR by injection of tunicamycin. Creld2 augments protein folding and creates an interlink between the UPR axes through its interaction with proteins involved in UPR. Thereby, Creld2 promotes tolerance to ER stress and recovery from acute stress. *Creld2*-deficiency leads to a dysregulated UPR, and causes the development of hepatic steatosis during ER stress conditions. Moreover, Creld2 enhancement of the UPR assists in the regulation of energy expenditure. Furthermore, we observed a sex dimorphism in humans with fatty liver disease, with only males showing an accumulation of CRELD2 protein in the liver. These results reveal a Creld2 function at the intersection between UPR and metabolic homeostasis and suggest a mechanism in which chronic ER stress underlies fatty liver disease in males.

## Introduction

The Cysteine-rich with EGF-like domains (Creld) protein family consists of two members – Creld1 and Creld2 – and is highly conserved across species [1,2]. The domain structure between these proteins is almost identical with the only difference of transmembrane domains being present in Creld1, but lacking in Creld2 [3]. This leads to the distinct localization and functions of these proteins within a cell: Creld1 is proposed to be anchored in the endoplasmic reticulum (ER) membrane with its domains facing the cytoplasm [4], although a plasma membrane bound form facing the extracellular matrix has been predicted as well [1]. We previously characterized the function of Creld1 being essential for heart development in mice [4]. Creld2 is localized predominantly within the ER, can be secreted [3], and has been characterized as a marker for kidney disease in urine [5] and for prosthetic joint infections in synovial fluid [6]. However, the physiological role of Creld2 *in vivo* remains elusive.

Several *in vitro* studies have identified *Creld2* as an ER-stress inducible gene, whose expression is dependent on the ER stress sensor activating transcription factor 6 (Atf6) activity [7,8]. Yet, for full gene expression induction the combined activity of all three ER stress sensors, namely inositol requiring enzyme 1 (Ire1), protein kinase RNA‐activated (PKR)‐like ER kinase (Perk) and Atf6, is required [9,10]. ER stress is characterized by an accumulation of mis- and/or unfolded proteins in the ER lumen. Consequently, cells activate the unfolded protein response (UPR), a network of signaling pathways that collectively aims at decreasing ER protein load by broad inhibition of protein synthesis and at the same time promotes the activation and production of proteins that increase the protein folding capacity and protein degradation. The latter include chaperones, shuttling proteins that promote secretion of proteins out of the ER and ER-associated protein degradation (ERAD) components [11,12].

The ER luminal domains of the ER stress sensors are bound and thereby inactivated by the chaperone Grp78 (glucose‐regulated protein 78, also known as heat shock protein A5 (Hspa5)). Upon ER stress Grp78 is sequestered by the accumulation of proteins in the ER lumen, causing the activation of the three sensors. Perk dimerizes and is activated by trans‐autophosphorylation of its kinase domains, leading to translational inhibition through eIF2α phosphorylation. This directly enhances the translation of DNA‐damage‐inducible 34 (Gadd34) and CAAT/enhancer‐binding protein (C/EBP) homologous protein (Chop). While Gadd34 serves as a feedback loop to dephosphorylate eIF2α allowing the cell to recover from translational inhibition, increased Chop expression may trigger cell death due to unresolved ER stress. The kinase Ire1 undergoes autophosphorylation resulting in splicing of the *X-box binding protein 1* (*sXbp1*) mRNA producing an active transcription factor, which induces the transcription of chaperones and ERAD pathway components. ER-membrane bound Atf6 is proteolytically cleaved and translocates to the nucleus where it activates target genes including ERAD components, chaperones, and major genes involved in lipid metabolism [13].

The liver is a highly secretory organ and serves frequently as the tissue to investigate ER stress. Previous studies demonstrate that induction of ER stress leads to hepatic steatosis through the direct and indirect regulation of lipid metabolism [14–16]. On the contrary, in genetic and dietary models of obesity, a link between hepatic stress and insulin resistance has been established [17–20], suggesting that the accumulation of lipids within hepatocytes and the resulting hepatocellular damage is linked to dysfunction of the ER. Therefore, the arrow of causality between hepatosteatosis and ER stress remains unclear.

Intriguingly, *Atf6* knockout mice do not exhibit developmental defects or any obvious phenotypes at steady state, yet show an abrogated response to ER stress under challenge with the ER stressor tunicamycin (Tm), which macroscopically manifests in the development of liver steatosis [15,21]. Further, Atf6 was shown to regulate physiological responses under diet-induced obesity [22], suggesting the UPR being upstream of hepatosteatosis.

To address the function of Creld2 within the Atf6 signaling pathway and its possible role in hepatic pathophysiology, we generated a *Creld2* knockout mouse and characterized its role during ER stress challenge and diet-induced obesity in the liver. We show that Creld2 deficiency promotes the development of liver steatosis when mice are burdened with ER stress, while lack of Creld2 ameliorates diet-induced hepatosteatosis. Further, we identify Creld2 protein interaction partners that are involved in ER-stress release, thereby placing Creld2 functionally into the cellular stress response of a cell *in vivo*. The conserved function of Creld2 across species is supported by the accumulation of CRELD2 in male patients with non-alcoholic steatohepatitis (NASH).

## Results

To investigate Creld2 function *in vivo* we generated a *Creld2-*knockout mouse model by replacing the endogenous locus with an enhanced green fluorescent protein (eGFP) (*Creld2*^*eGFP/eGFP*^, Figure 1A), allowing the use of both *Creld2*^*WT/eGFP*^ *and Creld2*^*eGFP/eGFP*^ as reporter mice for *Creld2* expression. We confirmed ubiquitous protein expression [23] as well as lack of Creld2 expression in *Creld2*^*eGFP/eGFP*^ mice in various tissues via Western blot (Figure 1B). *Creld2*^*eGFP/eGFP*^ mice were born in Mendelian ratios (Figure 1C) and were viable. In concordance with studies of *Atf6*^*-/-*^ mice, which do not show any anomalous phenotype during unchallenged conditions [15], young *Creld2*^*eGFP/eGFP*^ mice (1-2 month-old) did not show differences in body and liver weight (Figure 1D, E) or changes in tissue morphology as assessed histologically via hematoxylin-eosin (HE) and Masson-Goldner Trichrome (MGT) stain (Figure 1F).

**Figure 1:**
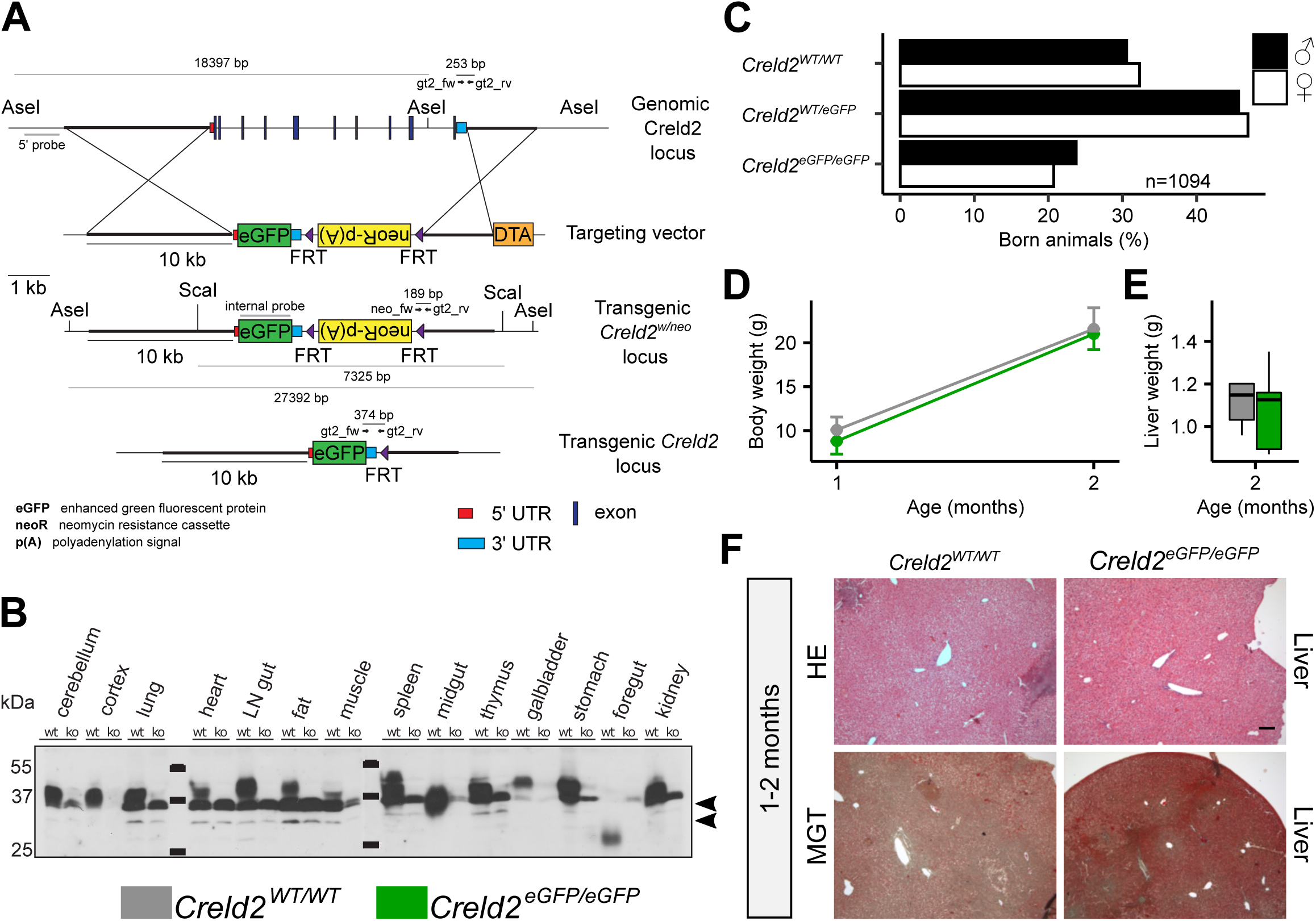
Generation and phenotypic analysis of the Creld2eGFP/eGFP mouse. **(A)** Gene targeting strategy. The complete genetic locus of Creld2 was replaced by an eGFP reporter via homologous recombination. The targeting vector comprised a neoR cassette flanked by FRT sites for positive ES cell selection. Following recombination, the neoR cassette was excised by FLP-mediated deletion. Primers for ES cell screening (gt2_fw, gt2_rv, neo_fw) and probes used for Southern blotting (5’ probe, internal probe) and the resulting fragment lengths (bp) are indicated. **(B)** Tissue expression profile of Creld2 and testing of the anti-Creld2 antibody specificity by immunoblotting. Arrowheads indicate unspecific protein bands. **(C)** Animals born from Creld2 WT/GFP x Creld2 WT/GFP matings. **(D)** Body weight of males (1 month: n = 5-7 and 2 months: n = 44-50). Error bars represent ± SD. **(E)** Liver weight of males (2 months: n = 5). Unpaired two-tailed t-test. **(F)** Histological analysis of liver by hematoxylin-eosin (HE) and Masson Goldner trichrome (MGT) in young Creld2eGFP/eGFP and wildtype mice. Representative for n = 8-10.

Given that *Creld2* is induced by Atf6 and that Atf6 was shown to regulate responses under high-fat diet induced obesity [22], we asked whether Creld2 is involved in regulating metabolic responses due to prolonged accumulation of lipids in hepatocytes. To induce the development of fatty liver, we placed 8-week-old *Creld2*^*eGFP/eGFP*^ males and wildtype littermates on control diet (CD) or high-fat diet (HFD). To test whether animals could recover from hepatosteatosis, we split the HFD group after 8 weeks with one group remaining on HFD for 4 more weeks and one group being switched to CD (HFD>CD, Figure 2A). Measuring mice body weights, we found that *Creld2*^*eGFP/eGFP*^ animals on CD gained less weight over time (Figure 2B). Similarly, after 8-12 weeks on HFD *Creld2*^*eGFP/eGFP*^ showed reduced body weights when compared to littermate controls (Figure 2B), but developed insulin resistance as measured by the homeostatic model assessment indices of insulin resistance (HOMA-IR) after 8 weeks (Figure 2C). Moreover, both genotypes increased the total tissue weight of livers and epididymal white adipose tissue (eWAT) (Figure S1A) indicating that *Creld2*^*eGFP/eGFP*^ animals were not protected from diet-induced obesity.

**Figure 2:**
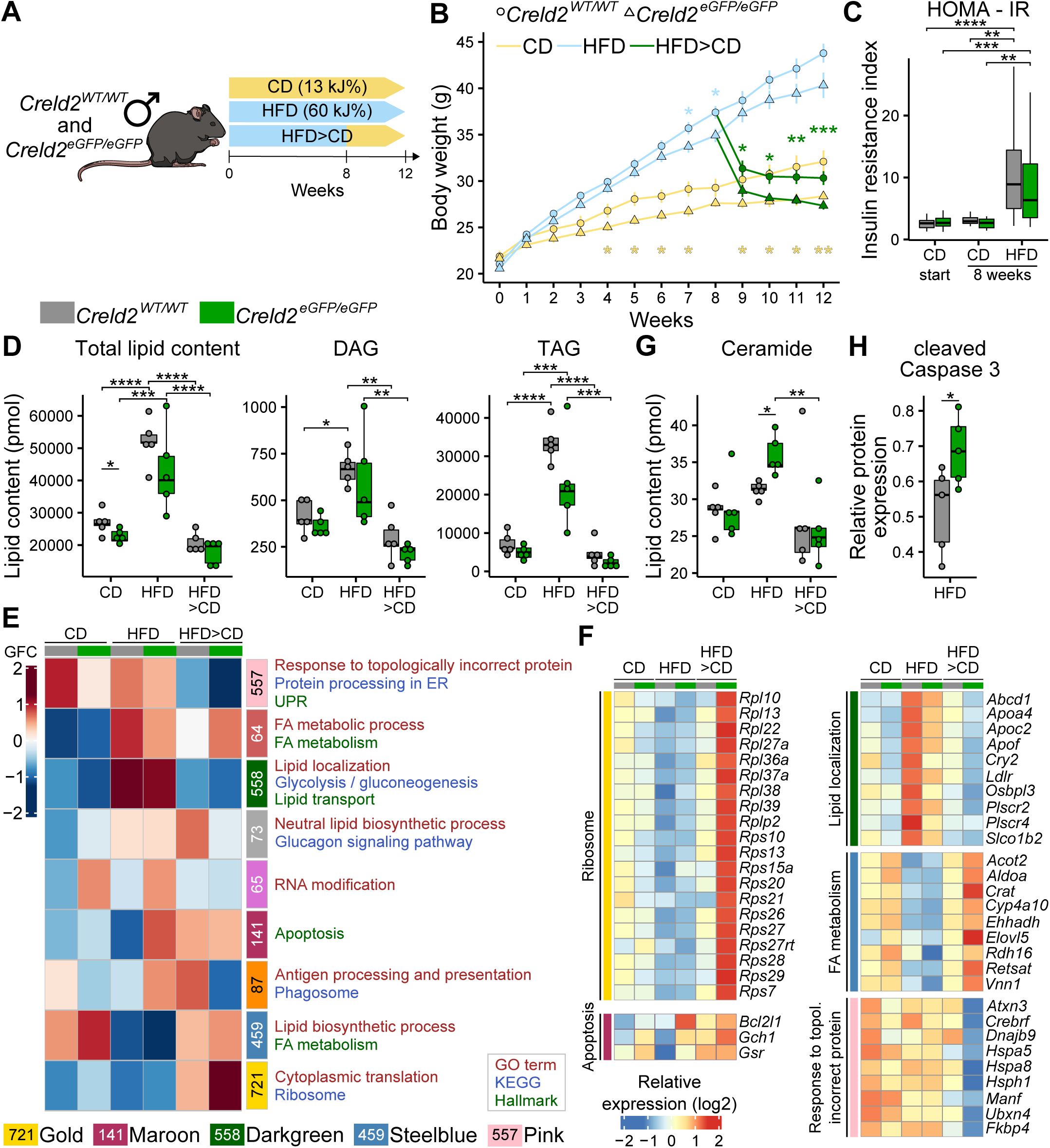
Tissue and molecular response of Creld2eGFP/eGFP mice to high-fat diet. **(A)** Experimental setup. **(B)** Body weight of male mice fed with CD (control diet; n = 13-14) and HFD (high-fat diet; n = 12-31) for 12 weeks, or HFD for 8 weeks following CD for 4 weeks (n = 14-19). **(C)** Analysis of insulin resistance induction by HFD via HOMA-IR (homeostatic model assessment of insulin resistance, CD start: n = 22-29; CD 8 weeks: n = 7-10; HDF 8 weeks: n = 17-20). One-way ANOVA with Tukey post-hoc test. **(D)** Lipid mass spectrometry of livers (n = 5 per condition). DAG: Diacylglycerol. TAG: Triacylglycerol. Circles represent individual mice. One-way ANOVA. **(E)** Co-expression network analysis (CoCena) performed on RNA-seq data of livers (n = 5 per condition). Numbers indicate genes belonging to a module. Colours represent cluster names: pink: 557 genes, indianred: 64 genes, darkgreen: 558 genes, darkgrey: 73 genes, orchid: 65 genes, maroon: 141 genes, darkorange: 87 genes, steelblue: 459 genes, gold: 721 genes. GFC: group fold change. **(F)** Heatmap representation of selected genes from (E), values are displayed as z scores. **(G)** Lipid mass spectrometry analysis for ceramides. Circles represent individual mice. One-way ANOVA with Tukey post-hoc test. **(H)** Western blot analysis for cleaved Caspase 3 (n = 5). Circles represent individual mice. Unpaired two-tailed t-test. * p < 0.05; ** p < 0.01; *** p < 0.001, **** p < 0.0001.

Histological assessment of the liver showed an increased oil-red-O (ORO) staining in HFD-fed mice (Figure S1B, C), indicating increased neutral lipid storage. This was accompanied by a decrease in PAS signal (Figure S1D, E) confirming diet-induced hepatosteatosis in both genotypes. Accumulation of lipids in general, which resulted mainly from TAG and DAG accumulation, was confirmed by shotgun lipidomics (Figure 2D). *Creld2*^*eGFP/eGFP*^ livers contained generally less lipids (Figure 2D), which could account for the on average lower liver weight (Figure S1A). HFD did not induce a fibrotic phenotype, nor did it affect liver function as assessed by MGT staining (Figure S1F, G) and alanine aminotransferase (ALT) concentrations in the serum (Figure S1H), respectively. When the HFD groups where switched to CD, their body, liver and eWAT weight returned to the level of animals that were kept on CD (Figure 2B, S1A). Further, ORO and PAS signals as well as the amount of lipids returned to baseline levels in both genotypes (Figure 2D, S1B-E). Together, our data indicate that on organismal and tissue level *Creld2-*deficient mice responded similarly to wildtype mice upon diet-induced obesity and lipid uptake as well as mobilization of lipids after a diet switch from HFD to CD.

To test whether lipid accumulation or subsequent lipid mobilization triggered differential cellular stress responses in *Creld2*^*eGFP/eGFP*^ and littermate control livers, we performed RNA-sequencing (RNA-seq) after 12 weeks on diet (Figure 2E, F, S1I). We made use of Construction of Co-expression network analysis 2 (CoCena^2^) to cluster the top 5000 variable genes into distinct modules based on the expression patterns across the two genotypes and three conditions (Figure 2E).

CoCena identified the gold module (721 genes) with globally upregulated genes in *Creld2*^*eGFP/eGFP*^ livers showing highest differences after HFD>CD diet (Figure 2E). Gene Ontology (GO) and KEGG enrichment analyses indicated an enrichment for cytoplasmic translation as well as ribosome proteins and ribosome production (Figure 2F). Additionally, the maroon module that showed an induction of 141 genes in *Creld2*^*eGFP/eGFP*^ livers after HFD returned ‘apoptosis’ as Hallmark term (Figure 2E) including *Bcl2l1, Gch1 and Gsr* (Figure 2F), suggesting that knockout livers may experience more apoptosis than wildtype controls. This notion was supported by an increase of ceramides in *Creld2*^*eGFP/eGFP*^ livers after HFD (Figure 2G), as their accumulation has been associated to apoptosis in hepatosteatosis [24]. Indeed, immunoblotting against cleaved Caspase3 confirmed that *Creld2*^*eGFP/eGFP*^ livers after HFD underwent increased apoptosis when compared to *Creld2*^*WT/WT*^ (Figure 2H, S1J).

Further, CoCena identified the darkgreen module, which contains genes being highly expressed after HFD diet, but which are globally downregulated in *Creld2*^*eGFP/eGFP*^ livers when comparing them to *Creld2*^*WT/WT*^ in each condition (Figure 2E, Table S1). GO and Hallmark terms indicated that many of the 558 genes were involved in the regulation of lipid localization and transport. Among these genes were Apoliporotein genes (*Apoa4, Apoc2, Apof*), which are important for lipid transport, genes involved in lipid distribution (*Abcd1, Ldlr, Osbpl3, Plscr2/4, Slco1b2*) and the circadian rhythm regulating transcription factor *Cry2*, which is responsible for the orchestration of physiological metabolism (Figure 2F).

The steelblue module (459 genes) includes genes, which are upregulated in *Creld2*^*eGFP/eGFP*^ animals during CD and HFD>CD recovery diets. Many of these genes are correlated to fatty acid (FA) metabolism, mostly involving genes for lipid catabolism (*Acot2, Ehhadh, Cyp4a10, Rdh16*) and some genes responsible for lipid anabolic processes (*Aldoa, Elovl5*) (Figure 2F), which may explain why *Creld2*^*eGFP/eGFP*^ animals and livers show reduced lipid amounts as they upregulate these genes even more than *Creld2*^*WT/WT*^ after HFD>CD recovery.

The pink module returned 557 genes that were downregulated in *Creld2*^*eGFP/eGFP*^ livers when compared to controls, particularly in the HFD>CD condition. These genes were involved in ‘responses to topologically incorrect protein’, ‘protein processing in the ER’ and the ‘UPR’ based on GO, KEGG and Hallmark enrichment analyses. Among these genes were chaperones and co-chaperones (*Dnajb9, Hspa8, Hsph1, Fkbp4*), including *Hspa5* (*Grp78*) (Figure 2F). We therefore investigated whether Grp78 expression and UPR activation were affected on the protein level. Yet, neither Grp78 nor phosphorylation levels of Perk and eIF2α were significantly changed in *Creld2*^*eGFP/eGFP*^ livers when compared to littermate controls, (Figure S1J, K). Further, the comparison of CD versus HFD groups per genotype did not reveal any sign of induced ER stress on transcript or protein level (Figure 2F, S1J-K), with Grp78 and Chop expression being even downregulated after HFD when compared to CD (Figure S1J-K). To test for other targets of the UPR that might be affected by the dietary change and that may have not been included in the CoCena^2^ analysis, we additionally performed a direct pairwise comparison of differentially expressed genes (lfc = 1.32; p-value = 0.05) between CD or HFD in *Creld2*^*WT/WT*^ or *Creld2*^*eGFP/eGFP*^ animals. None of these analyses returned genes involved in UPR (Table S2), indicating that 12 weeks of HFD do not induce ER stress in the liver.

Taken together, our data suggest that *Creld2*^*eGFP/eGFP*^ animals can generally cope with an excess of lipid uptake for a short time period of 12 weeks, despite transcriptional changes that hint towards altered lipid metabolic processes. Furthermore, neither transcriptome nor protein expression analysis revealed a lipid-driven ER stress response in the liver, despite an enhanced apoptotic phenotype in *Creld2*^*eGFP/eGFP*^mice livers.

Next, we turned to the question whether Creld2 is involved in the resolution of ER stress. To this end *Creld2*^*eGFP/eGFP*^ animals and littermate controls received a single intraperitoneal injection of the ER stressor Tm (1 mg/kg) or sucrose as control since Tm was prepared in a sucrose solution (Figure 3A). Two days post injection, mice were analyzed for the extent of fatty liver development, which served as surrogate readout of unresolved ER stress (Figure 3B, C). Histological analyses as well as shotgun lipidomics indicated that *Creld2*^*WT/WT*^ livers did not accumulate significant amounts of lipids or glycogen after Tm injection, while an increase of lipids after Tm injection could be observed in *Creld2*^*eGFP/eGFP*^ livers (Figure 3B, C, S2A-B). The slightly increased lipid storage in *Creld2*^*WT/WT*^ animals largely resulted from accumulation of cholesterol and cholesterol esters (Figure 3C). However, *Creld2*^*eGFP/eGFP*^ livers stored significantly more lipids than *Creld2*^*WT/WT*^ controls after Tm treatment, which could be mainly attributed to augmented TAG, DAG and cholesterol ester abundance (Figure 3B, C). This fatty liver phenotype was accompanied by a decreased PAS signal when compared to controls (Figure 3B, S2C). Further, MGT revealed an increased collagen abundance after Tm treatment in both genotypes (Figure S2D, E).

**Figure 3:**
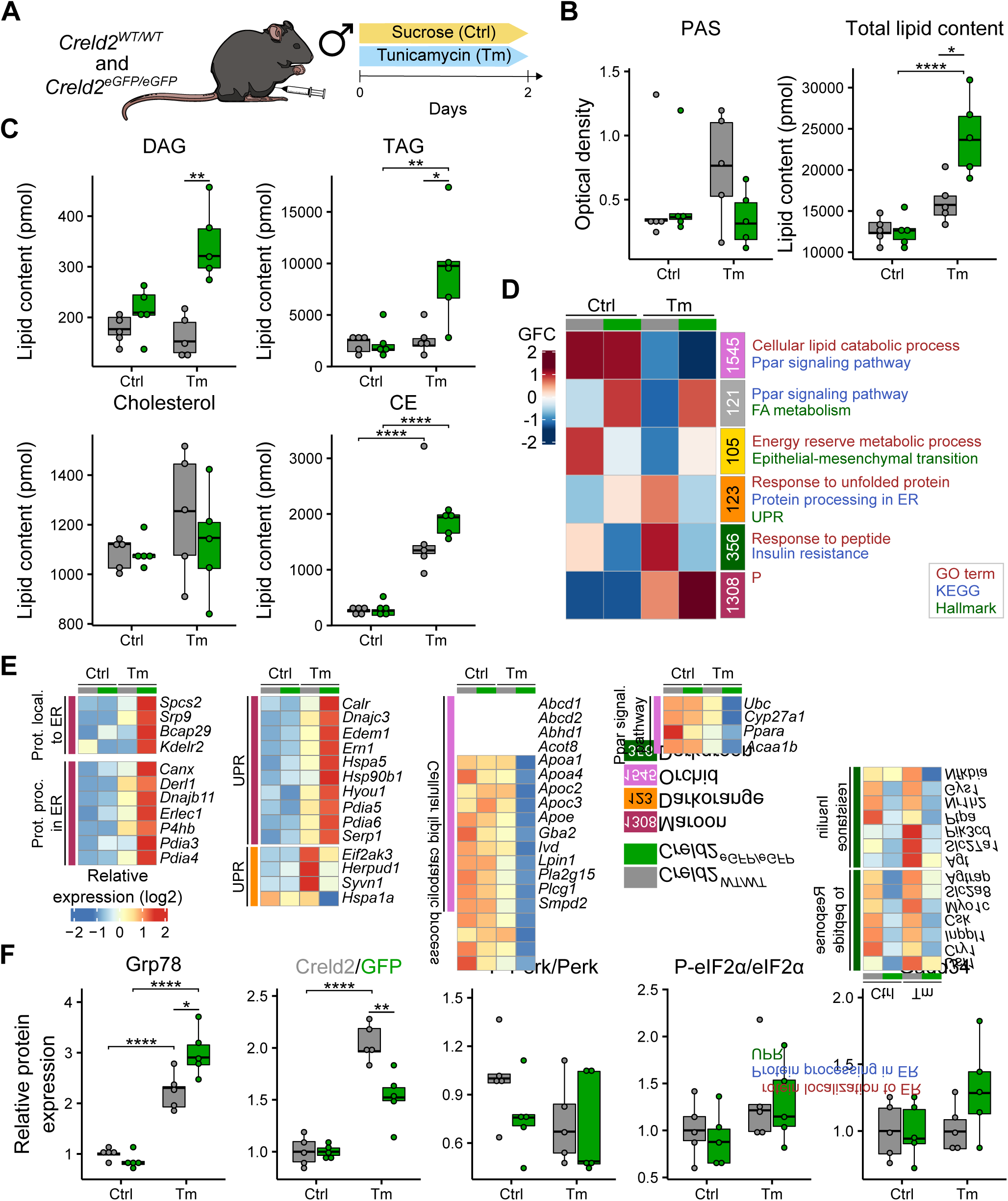
Tissue and molecular response of Creld2eGFP/eGFP mice to pharmacological ER stress induction. **(A)** Experimental setup. **(B)** Quantification of liver PAS stainings and mass spectrometrical assessment of liver lipids (n = 5 per condition). Circles represent individual mice. One-way ANOVA with Tukey post-hoc test. **(C)** Lipid mass spectrometry of livers (n = 5 per condition). DAG: Diacylglycerol. TAG: Triacylglycerol. CE: Cholesteryl esters. Circles represent individual mice. One-way ANOVA with Tukey post-hoc test. **(D)** Co-expression network analysis (CoCena) performed on RNA-seq data of livers (n = 5 per condition). Numbers indicate genes belonging to a module. Colours represent cluster names: orchid: 1545 genes, darkgrey: 121 genes, gold: 105 genes, darkorange: 123 genes, darkgreen: 356 genes, maroon: 1308 genes. GFC: group fold change. **(E)** Heatmap representation of selected genes from (D), values are displayed as z scores. **(F)** Western blot analysis for UPR components (n = 5 per condition). For Perk and eIF2α, ratios of phosphorylated (P-Perk and P-eIF2α) and unphosphorylated protein abundance are displayed. Circles represent individual mice. One-way ANOVA with Tukey post-hoc test. * p < 0.05; ** p < 0.01; *** p < 0.001, **** p < 0.0001.

To identify the molecular mechanisms that lead to augmented lipid accumulation in *Creld2*^*eGFP/eGFP*^ mice after ER-stress induction with Tm we performed RNA-seq on livers and analyzed the gene expression using principle component analysis (PCA, Figure S2F) and CoCena^2^ (Figure 3D, Table S3). The maroon module (1308 genes) contains genes that were upregulated after Tm treatment in both genotypes when compared to untreated mice, however, with even a higher expression in *Creld2*^*eGFP/eGFP*^ livers. GO, KEGG and Hallmark enrichment analyses returned terms such as ‘protein localization to ER’, ‘protein processing in ER’ and ‘UPR’ (Figure 3D), which comprised UPR-(*Hspa5, Hsp90b1, Ern1*) and ERAD-related genes (*Derl1, Edem1, Pdia3, Pdia4, Erlec1*) (Figure 3E). Tm treatment prevents protein glycosylation, which in turn induces ER stress and lipid accumulation [14] (Figure 3B, C). Wildtype livers largely recover from Tm-induced lipid perturbations after 2-3 days while keeping part of the UPR active, e.g. expression of chaperones Grp78 and Grp94 (*Hsp90b1*) [14,21]. Our data validate these observations showing that UPR is active after 2 days of Tm treatment and further suggest that *Creld2*^*eGFP/eGFP*^ livers undergo elevated ER stress when compared to *Creld2*^*WT/WT*^ controls. In contrast, the darkorange module (123 genes) identified genes that are downregulated in *Creld2*^*eGFP/eGFP*^ livers, and that also belong to the UPR (e.g. *Perk*/*Eif2ak3*) (Figure 3D, E). These two CoCena^2^ modules suggest that *Creld2*^*eGFP/eGFP*^ mice are only partly able to induce the UPR under ER stress, and, therefore, are incapable to properly resolve ongoing ER stress. This in turn results in increased hepatic lipid storage, supporting the active role of Creld2 during UPR and subsequent ER protein processing capacity.

In line with this hypothesis, the module orchid (1545 genes) classified genes that are downregulated after Tm treatment in both genotypes, with even stronger downregulation in *Creld2*^*eGFP/eGFP*^ livers (Figure 3D). GO and KEGG enrichment analysis of genes in the orchid module included mainly terms related to ‘cellular lipid catabolic process’ as well as ‘Ppar signaling pathway’ (Figure 3D), indicating that *Creld2*-deficient livers store increased amount of lipids, due to insufficient lipid catabolism (*Abcd1, Abcd2, Abhd1, Acot8, Gba2, Ivd, Pla2g15, Plcg1, Ppara, Acaa1b*) and diminished lipid transport (*Apoa1, Apoa4, Apoc2, Apoc3, Apoe*) (Figure 3E).

The module darkgreen identified 356 genes that were globally downregulated in *Creld2*^*eGFP/eGFP*^ livers, with highest differences after Tm treatment (Figure 3D). These genes enriched largely for ‘response to peptide’ and ‘insulin resistance’ as determined by GO and KEGG pathway enrichment analyses. Among enzymes regulating energy homeostasis were *Slc27a1, Gys1* and *Nr1h2* (insulin resistance) as well as *Usf1, Cry1* and *Slc2a8* (response to peptide), which collectively regulate lipid and glucose metabolism. In addition, expression of genes associated with signal transduction (*Pik3cd, Nfkbia*) and regulation of cellular growth (*Ptpa, Inppl1, Csk*) is disrupted in *Creld2*^*eGFP/eGFP*^ livers, exhibiting further aspects of Creld2-dependent regulatory processes (Figure 3E). Summarized, CoCena^2^ analyses of Tm challenged livers reveal a Creld2-dependent dysregulation of genes important for the resolution of ER stress resulting in altered lipid and glucose metabolism, and cellular signaling processes in *Creld2*-deficient mice livers, which subsequently leads to increased lipid storage.

To test this hypothesis, we first analyzed Grp78 protein abundance after Tm treatment as an indicator for protein accumulation in the ER lumen. Similar to RNA-seq results, Grp78 was upregulated in wildtype animals after Tm when compared to control treatment, but was even more abundant in *Creld2*^*eGFP/eGFP*^ livers (Figure 3F, S2G). Of note, both GFP expression in *Creld2*^*eGFP/eGFP*^ and Creld2 expression in *Creld2*^*WT/WT*^ littermate controls was increased upon both treatment conditions (Figure 3F, S2G), rendering Creld2 expression and the *Creld2*^*WT/eGFP*^ mouse as an efficient tool to monitor cellular stress. We next tested the activation of the Perk-axis via analysis of Perk and eIF2α phosphorylation and Gadd34 protein abundance. Despite the increased levels of Gpr78, we did not detect enhanced activation of the Perk-axis in *Creld2*^*eGFP/eGFP*^ livers (Figure 3F, S2G). In summary, induction of ER stress via Tm causes exacerbated ER stress in *Creld2*^*eGFP/eGFP*^ mice, as indicated by increased Grp78 expression and consequently augmented hepatic lipid storage, supporting the notion that Creld2 is required to maintain tissue homeostasis in order to prevent lipid accumulation due to long-lasting and unresolved ER stress.

To investigate the molecular function of Creld2 in UPR and metabolic homeostasis we performed co-immunoprecipitation experiments by tandem affinity purification [25] and subsequent mass spectrometry. We used murine Creld2 tagged either at the N- or C-terminus to exclude inhibition of protein binding due to the fused Strep-Flag tag. Moreover, Creld2 constructs were either transiently expressed (72 h) or stably integrated into the genome of HEK239 cells (Figure 4A) to *i)* enable the characterization of specific interactions with chaperones, since a strong transient overexpression increases the unspecific co-purification of heat shock proteins [25] and *ii)* select for highly conserved interactions, which would also occur in this inter-species (mouse-human) setup. Using a stringent filtering pipeline and imputation for missing values (Figure 4B, see also methods), we found 31 putative interaction partners (Figure 4C, Table S4). Analysis for GO enrichment returned terms such as ‘response to ER stress’, ‘protein folding’, and ‘cell redox homeostasis’ (Figure 4D). Many chaperones and protein disulphide-isomerases (PDI) were present in the transient condition (C2-SF trans), some of which (HSPA8, HSP90B, HYOU1) have been previously proposed to interact with Creld2 [26]. However, since these proteins were not enriched in the stably transfected cells (FS-C2 stab and C2-SF stab) we propose that these represent unspecific co-purification products [25] underlining the constraints of co-immunoprecipitation assays in transiently transfected cell lines.

**Figure 4:**
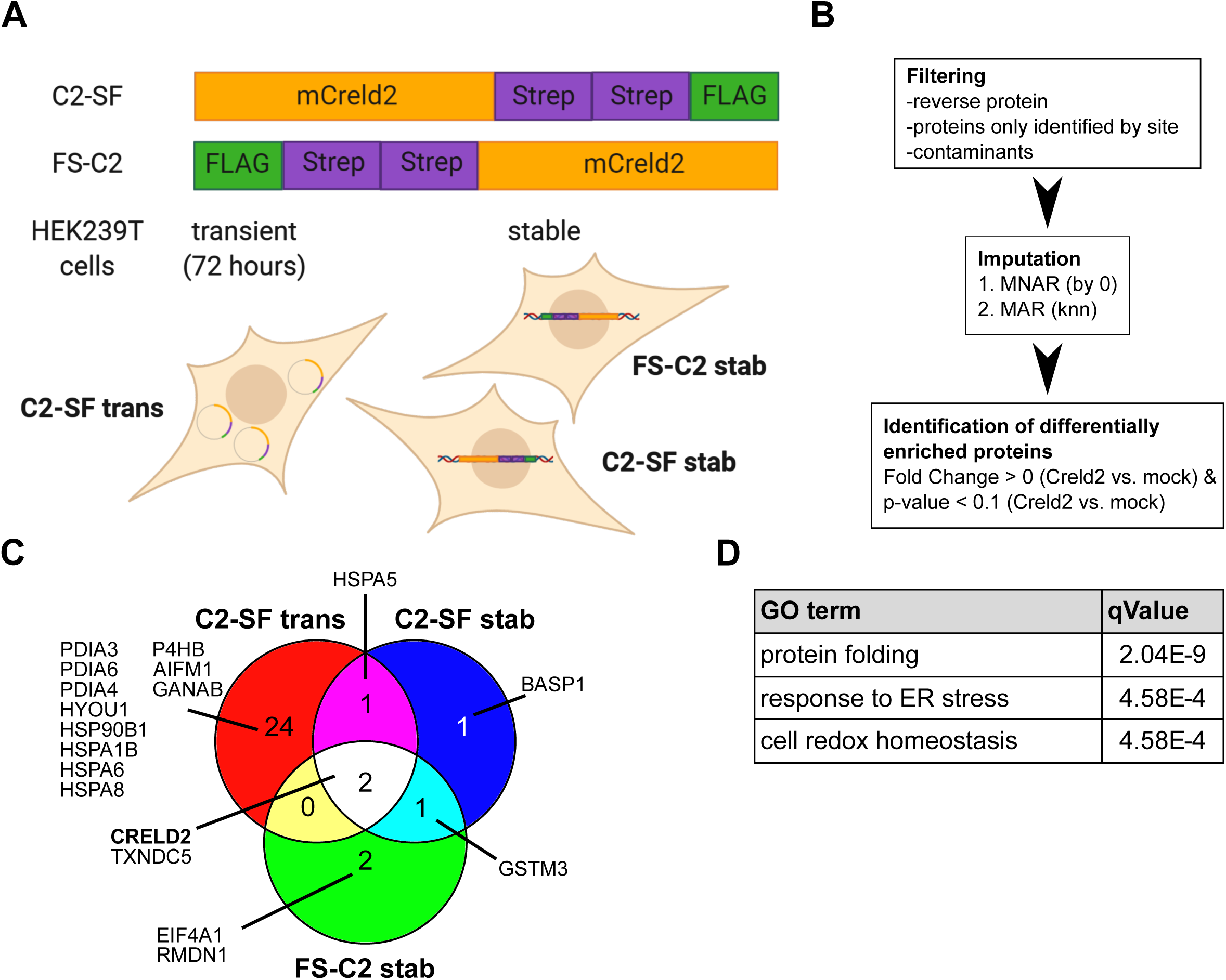
Creld2 interacts with proteins involved in protein folding, response to ER stress and detoxification. **(A)** Schematic representation of constructs and expression conditions used for tandem-affinity-purification of tagged murine Creld2. **(B)** Workflow chart for analysis of mass spectrometry data. MNAR values in the dataset were first imputed by 0, followed MAR imputation by knn algorithm. MNAR: missing not at random; MAR: missing at random; knn: k-nearest neighbours. **(C)** Enriched Creld2 interaction partners across the different expression conditions. **(D)** Top three enriched GO terms resulting from enrichment analysis of all co-purified proteins.

We thus focused our analysis on candidates present in at least 2 conditions. This led to the identification of Creld2 itself, as expected, as well as HSPA5 (GRP78), thioredoxin domain-containing 5 (TXNDC5) and glutathione S-transferase Mu3 (GSTM3) as potent specific Creld2 binding partners (Figure 4C). Interaction with GRP78 [26] further endorses an active role of Creld2 in regulating UPR, as it may sequester the chaperone away from Perk/Atf6/Ire1, thereby, promoting the activation of the three UPR axes. TXNDC5, also known as endo PDI, is a thioredoxin peroxidase whose expression is regulated by ATF6 and sXBP1. Functionally it is proposed to reduce misfolded protein load [27]. Intriguingly, also Creld2 has been proposed to have PDI-like functions [26], hinting towards a synergistic mechanism between these proteins to resolve ER stress. GSTM3 is a member of the glutathione S-transferases superfamily that promotes detoxification of reactive oxygen species (ROS) [28]. Oxidative stress and UPR are tightly interlinked, exemplified by PERK being upstream of the transcription factor NRF2 that counteracts the toxic effects of ROS produced upon ER stress during protein folding, e.g. via PDI activity [29]. Thus, the interaction of Creld2 with GSTM3 may represent an additional point of crosstalk between these cellular stresses. In summary, our data strengthen the hypothesis of an active involvement of Creld2 in UPR via interaction with chaperones and enzymes, whose activity is required to overcome cellular stress.

Overall, our data suggested that Creld2 function is required to maintain hepatic homeostasis under ER stress responses in mice. Therefore, we next asked whether human CRELD2 would play a role under pathophysiological conditions in the liver and analysed a cohort of 57 obese patients with varying severity of non-alcoholic fatty liver disease (NAFLD) and 4 controls for CRELD2 expression. We first divided patients into two groups - with or without non-alcoholic steatohepatitis (NASH) as diagnosed using the SAF (Steatosis, Activity and Fibrosis) score, which increases with disease progression [30]. Intriguingly, both CRELD2 transcript and protein levels in the liver were significantly upregulated in male patients with NASH, while expression in female patients remained unaffected (Figure 5A). Control patients showed similar expression to NAFL patients. To test whether secreted CRELD2 could be used as a diagnostic marker for NAFLD severity we plotted the serum concentration of CRELD2 against the SAF score (Figure 5B). While no correlation was observed in female patients, males had an inverse correlation with low levels of CRELD2 at higher SAF scores, which may hint towards a retention mechanism for CRELD2 in the tissue during hepatic pathophysiology. In summary, CRELD2 shows a sex-specific function during NAFLD in humans with an intra-hepatic upregulation of CRELD2 expression, which seems to be associated with progression to NASH in male patients.

**Figure 5:**
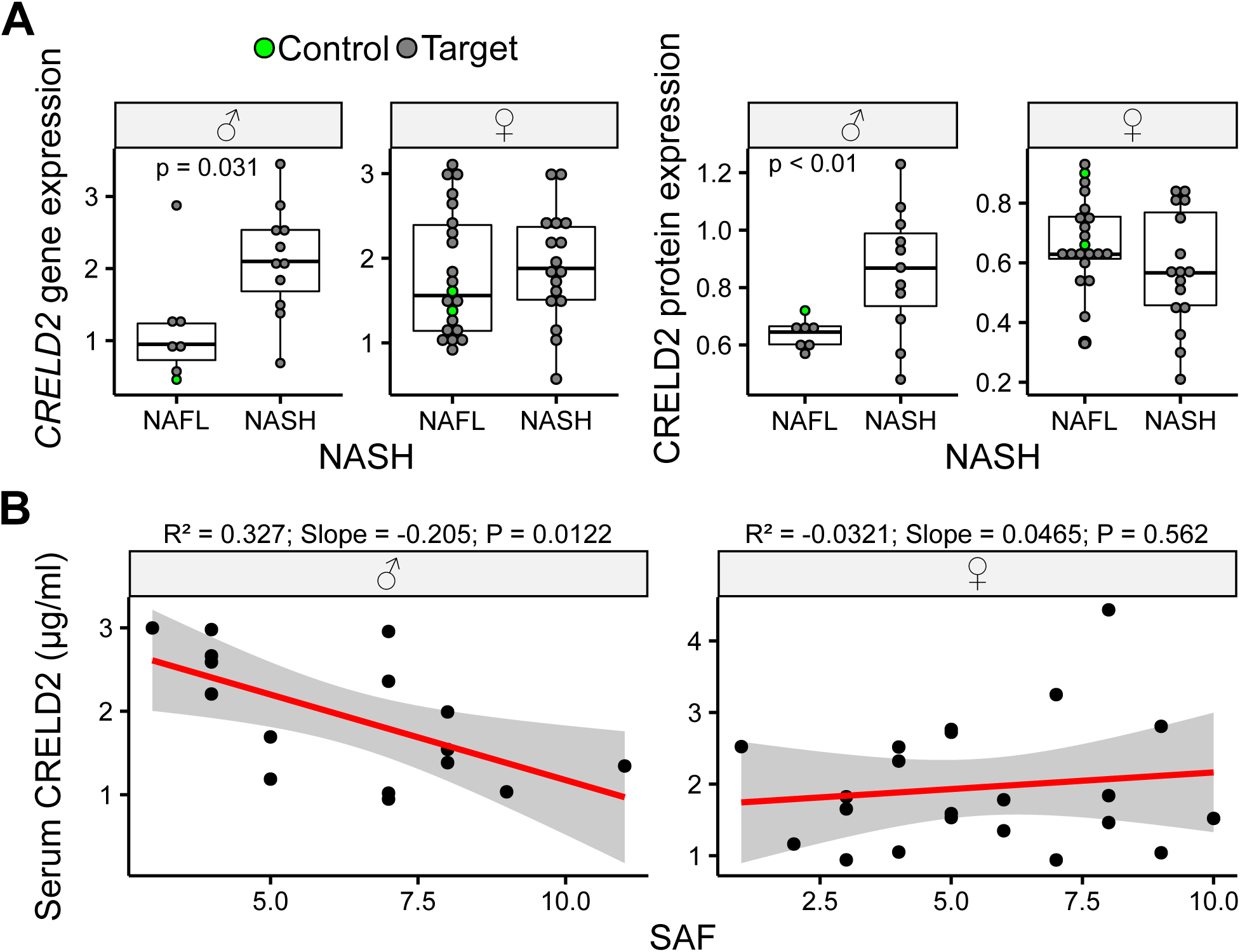
Hepatic CRELD2 expression increases in patients with non-alcoholic steatohepatitis. **(A)** Hepatic gene and protein expression levels of CRELD2 in male (♂) and female (♀) non-alcoholic fatty liver disease (NAFLD) patients (grey circles) with or without NASH (non-alcoholic steatohepatitis) and healthy controls (green circles). Cohort is comprised of 57 patients. Circles represent individual patients. Unpaired two-tailed t-test. **(B)** Analysis of CRELD2 serum concertation in male (♂) and female (♀) patients plotted against the Steatosis Activity and Fibrosis (SAF) score (cohort comprised of 40 patients). Linear regression model.

## Discussion

Our study shows that Creld2 plays an active role during UPR and contributes to the overall maintenance of cell homeostasis and resolution of ER stress. Using the newly generated *Creld2*^*eGFP/eGFP*^ transgenic mouse model, we characterized the hepatic stress response of *Creld2*-deficient mice. *Creld2*^*eGFP/eGFP*^ mice do not show gross phenotypic differences during steady state compared to wildtype controls. However, after challenge with Tm, *Creld2*-deficient livers show perturbated induction of UPR, which ultimately leads to dysregulated lipid and glucose metabolism resulting in exacerbated development of NAFLD. These findings are congruent with previous reports about *Atf6* knockout mice [15,21], suggesting the involvement of Creld2 in regulating a proper response to ER stress downstream of Atf6. This is further evidenced by increased susceptibility of CRELD2-deficient Neuro2a cells to treatment with Tm [23].

Dysregulation of the UPR in *Creld2*-deficient livers is evident by increased expression of major UPR components, e.g. *Edem1, Ern1* (*Ire1*) and *Hspa5*. At the same time they show decreased expression of *Perk* and Perk downstream target genes, such as *Herpud1* and *Hspa1a* [10,31], as well as *Syvn1*, which is induced by sXbp1 possibly in combined action with Atf6 [32]. UPR dysregulation is accompanied by downregulated gene expression of important components for lipid transport and catabolism, energy metabolism and insulin signalling, which reportedly come along with an impaired ER stress response [14,33–35]. Thus, *Creld2*-deficient livers show overactive UPR but are unable to fully activate the gene expression programs of all three UPR branches. Since Creld2 expression is induced by the Atf6 pathway we hypothesized that Creld2 is involved in the regulation of the Atf6 axis. In fact, a majority of highly induced UPR genes in *Creld2*-deficient livers are Atf6 target genes (*Calr, Pdia4, Pdia6, Hsp90b1, Hyou1)* [10]. Yet, Creld2^*eGFP/eGFP*^ mice also display deficient induction of parts of the Perk and Ire1 pathway on transcriptional level, suggesting that Creld2 is involved in a cross-talk between the three UPR axes.

In contrast to pharmacological ER stress challenge, *Creld2*^*eGFP/eGFP*^ mice fed with HFD display an ameliorated dietary induced obesity phenotype, hinting towards another layer of Creld2 impact on organ function. In this model, *Creld2*-deficient mice reveal lower body weights and reduced liver steatosis. These findings are accompanied by downregulated expression of genes important for lipid localization and transport but increased expression of genes necessary for lipid catabolism, a process required for energy production from fatty acids. These observations indicate that Creld2^*eGFP/eGFP*^ mice have a fundamentally higher energy expenditure. The underlying cause for the higher energy demand might be the upregulated expression of ribosomal genes, especially during HFD recovery (HFD>CD). Since the production of ribosomal proteins is a highly energy consuming process [36], *Creld2*-deficient mice may upregulate ATP synthesis and, therefore, increase the utility and catabolism of lipids leading to the lower body weights and less accumulation of lipids in their livers as compared to littermate controls. Concomitantly, livers of Creld2^*eGFP/eGFP*^ animals display reduced gene expression of UPR components in all conditions with lowest induction during HFD recovery. These observations can be linked to previous reports showing that perturbation of Perk downstream components eIF2α [37] or Atf4 [38], lead to diminished hepatosteatosis during high-fat diet and leaner mice exhibiting increased energy expenditure, respectively. In addition, Perk inhibits the expression of ribosomal genes [31], hinting towards an impairment of Perk downstream targets in *Creld2*-deficient mice, which results in the upregulation of ribosomal genes followed by increased fatty acid catabolism for ATP production due to the increased energy demand in these mice.

Furthermore, we found no evidence that lipid accumulation through HFD *per se* induces UPR in wildtype or *Creld2*^*eGFP/eGFP*^ livers. Yet, livers from *Creld2*^*eGFP/eGFP*^ animals show first signs of cellular stress and apoptosis when compared to controls, which underlines the UPR-independent protective function of Creld2 upon oxidative stress and ceramide-mediated hepatotoxicity caused by HFD [39,40]. Further, the diet switch from HFD to CD did not promote ER-stress induction in both genotypes, but rather reduced the expression of chaperones in *Creld2*^*eGFP/eGFP*^ livers. Thus, we conclude that UPR does not play a major role during diet-induced hepatic steatosis as long as this pathophysiology is reversible and did not progress to NASH. Taken together, our data support the hypothesis that Creld2 is required to protect cells against ongoing stress, although the cells retain the ability to partially compensate through other signalling pathways.

Analysis of Creld2 interaction partners further implicates an active role in regulating the UPR: Creld2 protein accumulates 48 h after induced ER stress and has the capacity to bind to Grp78. This in turn, may have two effects *i*) Grp78 would be sequestered from Perk/Ire1/Atf6 and thereby these sensors would be activated or *ii*) Creld2 might act as a co-chaperone and aid Grp78 in protein folding. Further, Creld2 binds to the endo PDI Txndc5 and glutathione S-transferase Gstm3, two proteins important for ER stress reduction [41,42] and ROS detoxification [43], respectively. Txndc5 was reported to be transcriptionally regulated by Atf6 and sXbp1 [27], which places Creld2 function at the intersection between protein folding processes and the three UPR axes. How binding to Creld2 influences the function of these proteins remains to be investigated. Nevertheless, based on our mouse work we propose that Creld2 has a dual role during steady state and cellular stress conditions: it maintains balanced basal and ER stress responsive UPR activation, probably by modulating the Perk- and Atf6-dependent axes through binding to Grp78, and at the same time it contributes to the reduction of misfolded proteins either through its own putative PDI activity or by promoting the PDI activity of Txndc5 and detoxification function of Gstm3.

In humans, the fundamental causes for NAFLD development can be of various nature. However, the progression to NASH and hepatocellular carcinoma (HCC) is associated with or driven by increased and unresolved ER stress [44,45]. Our CRELD2 expression analyses in livers of human NAFLD patients provide evidence for the importance of CRELD2 in the amelioration and reversion of NASH. Intriguingly, increased liver CRELD2 levels and the negative correlation of serum CRELD2 with disease severity could only be observed in male patients, while females did not show any differences. Sex differences in NAFLD have been demonstrated in both, rodent models and in human disease, to be more prevalent in men and postmenopausal women compared to premenopausal women, which is mostly attributed to hormone homeostasis and visceral fat accumulation [46,47] (Figure 6). We therefore favor the hypothesis that the hepatic ER stress response is sex dependent and thus a risk factor for the development of NAFLD and progression to NASH or HCC particularly in males. This is in line with our results showing that CRELD2 accumulates only in livers of male NASH patients, as well as a previous study demonstrating that kidneys of male mice are more susceptible to ER stress-induced acute kidney injury than those of females due to a testosterone-dependent mechanism [48]. Further, CRELD2 - among six other proteins that maintain ER homeostasis - was identified as an adverse prognostic biomarker for overall survival in HCC patients [49] underlining that elevated and unresolved ER stress can promote liver pathophysiology. Therefore, it will be essential to take the sexual dimorphism into account in the future when studying the contribution of ER stress to hepatic steatosis as well as in the course of NAFLD treatment with UPR modulating agents.

**Figure 6:**
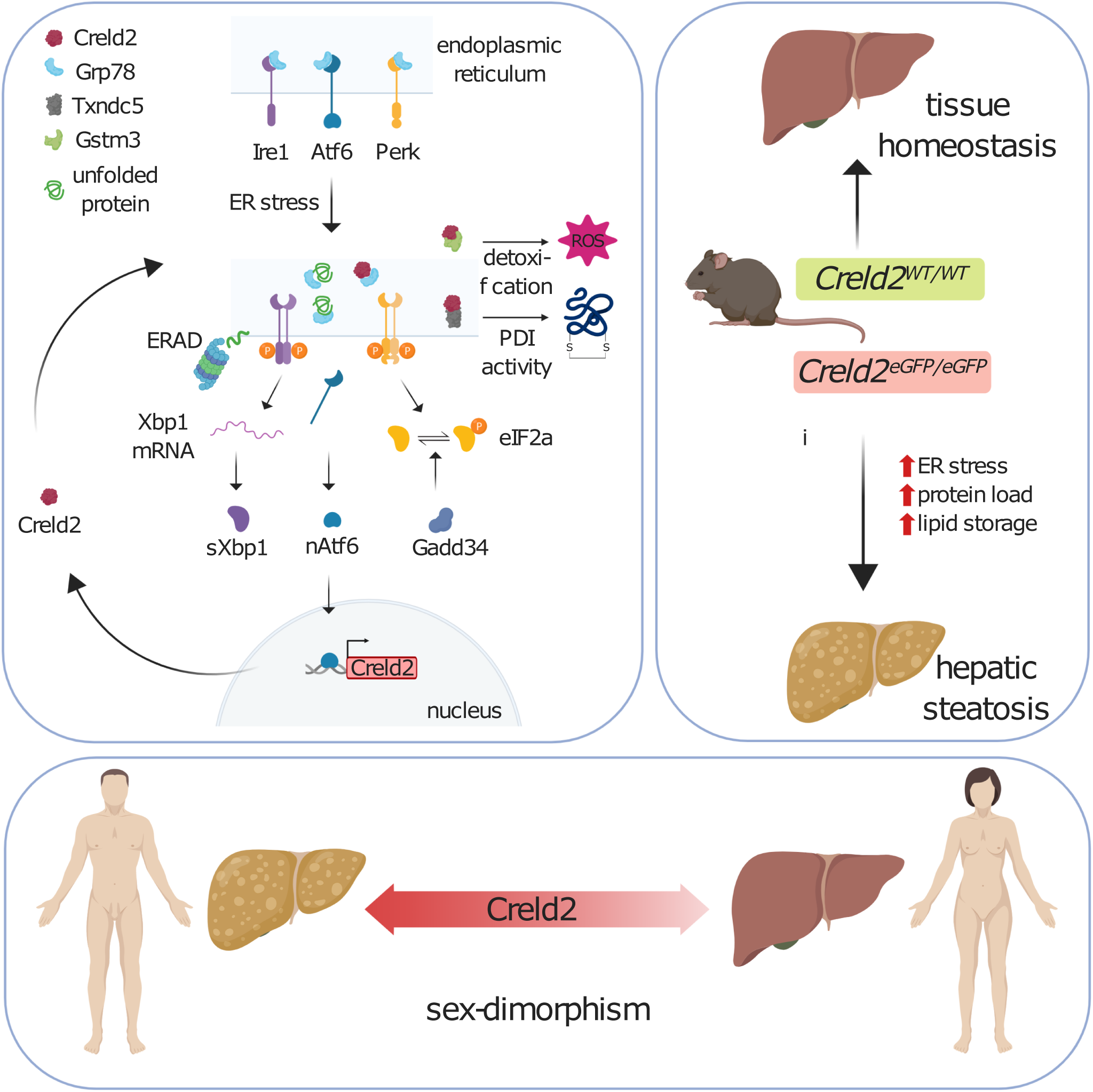
Graphical summary of Creld2 function. Summarizing schematic of Creld2 functionality, PDI: protein disulfide isomerase; ROS: reactive oxygen species.

## Materials and methods

### Mouse work

For the generation of a conventional *Creld2*^*eGFP/eGFP*^ mouse the open reading frame (ORF) of the *Creld2* gene was replaced by an enhanced green fluorescent protein (eGFP) and a neomycin resistance cassette (neoR) flanked by two FRT sites via homologous recombination. False positive ES cell clones were excluded by a diphtheria toxin A (DTA) cassette, which was introduced after the 3’ homologous region of the targeting vector. Positive ES cell clones were selected by PCR against the neoR cassette and confirmed by southern blot via an internal and 5’ external probe. Heterozygous Creld2^WT/eGFP/neo^ offspring from blastocyst injections of positive ES cell clones were crossed with mice ubiquitously expressing FLP recombinase for excision of the neoR cassette. neoR deletion was confirmed by PCR. Mice were provided with standard chow and autoclaved water ad libitum. Genotyping primers: Creld2 wt genotype fw 5’-CCTGAGCTGTCCTTAGAAAGTTGCTAG-3’, Creld2 ko genotype fw 5’-GCCCGACAACCACTACCTGAGC-3’, Creld2 genotype rev 5’-GGGGTTCATGTCCATGGGCCAC-3’.

To induce hepatic UPR, 8-9 weeks-old male mice were administered with Tm (1 mg/kg in 150 mM sucrose) or a vehicle control sucrose solution (150 mM) via i.p. injection and sacrificed after 48 h.

For HFD experiments, male mice received standard chow until 6 weeks of age. Afterwards, standard chow was replaced by a control diet (CD; metabolizable energy: 62 % from carbohydrates, 11 % from fat, 27 % from protein; ssniff EF D12450B* mod. LS) for 2 weeks prior to the beginning of the experiment. Subsequently, 8-week-old mice were either fed with CD or HFD (metabolizable energy: 22 % from carbohydrates, 54 % from fat, 24 % from protein; ssniff EF acc. D12492 (I) mod.) for 12 weeks or mice were kept on HFD for 8 weeks and CD for 4 weeks (HFD>CD). Mice were weighed once a week. For the determination of the Homeostatic model assessment of insulin resistance (HOMA-IR) [50], mice were fasted for 6 hours and blood was drawn from 8-week-old mice at the beginning of the feeding period and from mice being fed CD or HFD for 8 weeks. Blood samples were analysed for blood glucose using a glucose meter (Accu-Chek, Aviva) and blood serum insulin levels (Mouse Insulin Elisa kit, Thermo Scientific) according to the manufacturers’ instructions. HOMA-IR indices were calculated using the formula: (Glucose (mg/ml) x Insulin(µU/ml))/405.

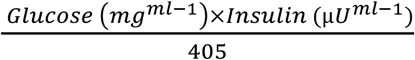

All mice were kept under standard SPF housing conditions according to the approved animal license 84-02.04.2014.A335 with a 12 h dark/light cycle. All animal experiments were approved by the government under the license 84-02.04.2017.A335

### Histological stainings

For paraffin stains, tissues were fixed in 4% PFA (Thermo Scientific). After fixation, tissues were dehydrated, paraffin embedded and cut at a thickness of 5 µm. Subsequently, sections were deparaffinized, rehydrated and stained with Hematoxylin followed by Eosin (VWR), PAS (Roth) or MGT (Merck) according to the manufacturer’s protocol. After dehydration, tissues were mounted in Entellan. For Cryo-embedded (Tissue-Tek, Sakura) tissue stains, sections were fixed in 4% PFA and stained with freshly prepared oil-red-O (Sigma-Aldrich) working solution according to the manufacturer’s instructions followed by mounting in Kaiser’s glycerol gelatine (Merck). Quantitative analyses of PAS, MGT and ORO stainings were done using ImageJ2 software. Scripts for quantifications are provided upon reasonable request.

### qPCR

RNA was isolated using the Nucleospin RNA II kit (Macherey & Nagel) according to the manufacturer’s instructions and reverse transcribed into cDNA by using the QantiTect reverse transcription kit (Qiagen) according to the manufacturer’s instructions. For qRT-PCR reactions, primer pairs (see Table S5) were mixed with cDNA and iQ SYBR’Green Supermix (Bio-Rad), and qRT-PCR was run on an iQs Real-Time PCR Detection System (Bio-Rad). *CRELD2* gene expression was normalized to the reference gene *HPRT* (hypoxanthine-guanine phosphoribosyltransferase). Relative gene expression was calculated from the threshold cycles in relation to the reference gene and controls without NAFLD. Reactions were performed on a CFX96 Touch qPCR System (Bio-Rad).

### Library preparation for RNA sequencing

mRNA was converted into libraries of double stranded cDNA molecules as a template for high throughput sequencing following the SMART-Seq2 protocol [51]. Shortly, mRNA was primed for SMART reverse transcription from 5 ng of total RNA using poly-T oligos. cDNA was pre-amplified by SMART ISPCR. Fragmentation was performed using the Illumina Nextera XT kit, followed by PCR amplification and indexing. Size-selection and purification of library fragments with preferentially 300-400 bp in length was performed using SPRIBeads (Beckman-Coulter). The size-distribution of cDNA libraries was measured using the Agilent high sensitivity D5000 assay on a Tapestation 4200 system (Agilent). cDNA libraries were quantified using a Qubit high sensitivity dsDNA assay. 75 bp single-end sequencing was performed on a NextSeq500 system using High Output v2.5 chemistry. Base calling from base call files, alignment to the Mus musculus reference genome mm10 from UCSC and file conversion to fastq files were achieved by Illumina standard pipeline scripts (STAR version, bcl2fastq2 v.2.20).

### RNA sequencing analysis

Kallisto pseudo-alignment [52] was used to quantify abundances of transcripts - read counts - from the bulk RNA-seq data. The kallisto read counts were used as input to DESeq2 [53] for calculation of normalized signal and differential gene expression. Genes are excluded from the analysis where the total read count of all samples is less than 10. This pre-filtering step removes genes in which there are very few reads, in order to reduce the memory size, and to increase the speed of the calculation. The diet and drug induced ER-stress dataset then comprised 25308 and 23985 genes respectively. DESeq Dataset (dds) was created to store the read counts and the intermediate estimated quantities with a merged design formula consist of genotype (*Creld2*^*WT/WT*^ or *Creld2*^*eGFP/eGFP*^) and condition (dietary or pharmacological). Regularized log transformations (rlog) were used for the downstream analysis (clustering). Rlog produces transformed data on the log2 scale which has been normalized with respect to library size. Unwanted surrogate variables were estimated and modelled via Surrogate Variable Analysis (SVA) [54] using the “leek” method. In the diet related experiment 4 surrogate variables (SV) were identified and modelled. For the pharmacologically-induced dataset 3 SVs were identified and modelled. For DE gene analysis of CD versus HFD conditions, lfc threshold was set to 1.32. and the p-value threshold to 0.05. The top 5000 variable genes were selected and used as an input for co-expressional network analysis using CoCena^2^ (https://github.com/UlasThomas/CoCena2). Pearson correlation analysis was performed with correlational significance measure p<0.05. For the diet-induced dataset 0.706 and for the pharmacologically-induced dataset 0.659 was identified as correlation coefficient cut-off. Based on a greedy approach, Louvain community detecting algorithm [55] was used to cluster the genes of the diet-induced dataset based on expression pattern between the different conditions. For the pharmacologically-induced dataset the infomap community detection algorithm [56] was used. Clustering was repeated 100 times. Minimal number of genes per cluster was set to 50. Gene set enrichment analysis (GSEA) was performed on all cluster genes using the following knowledge-bases: Hallmark, Reactome, GO, KEGG and DO. ClusterProfiler package [57] was used for all GSEAs using default options with the respective correlation coefficient cut-off for each dataset. The prior mechanisms associated with the Co-Cena clusters were manually selected from the GSEA results based on prior knowledge.

### Thin layer chromatography

Total lipids were extracted as described by Bligh and Dyer [58].Livers were homogenized in ice cold H^2^O (50mg wet weight/ml) in a Precellys homogenizer (Peqlab Biotechnology, Germany). 3 ml of chloroform/methanol 1:2 (v:v) were added to 800 µl homogenate and vortexed. 1 ml of chloroform was added to the mixture and vortexed prior to addition of 1 ml H^2^O and thoroughly vortexed. Mixture was centrifuged at 1000 g for 5 min at RT and the lower solvent phase was recovered. Same volumes of the solvent phase were evaporated under nitrogen and lipids were resuspended in 100 µl of chloroform/methanol 1:1 (v:v) (Merck). Lipid extracts corresponding to 1 mg wet liver weight were applied onto HPTLC silica gel 60 plates (10 × 20 cm: Merck) and plates were developed in *n*-hexane/diethylether/acetic acid 70:30:5 (v/v/v) (Merck, Roth) in a developing chamber (CAMAG, Switzerland). Dried plates were soaked in charring solution (copper sulfate (CuSO^4^) 10% and phosphoric acid (H^2^PO^4^) 8% (Roth)) and heated to 180 °C for 5-10 min for lipid visualization.

### Lipid mass spectrometry

Livers were homogenized in ddH^2^O at a concentration of 20 mg (wet weight)/ml using a Precellys homogenizer (Peqlab Biotechnolog). For lipid extraction, 50 µl of the homogenate were added to 500 µl of extraction mix (CHCl^3^/MeOH 1/5 containing internal standards: 210 pmol PE(31:1), 396 pmol PC(31:1), 98 pmol PS(31:1), 84 pmol PI(34:0), 56 pmol PA(31:1), 51 pmol PG (28:0), 28 pmol CL(56:0), 39 pmol LPA (17:0), 35 pmol LPC(17:1), 38 pmol LPE (17:0), 32 pmol Cer(17:0), 99 pmol SM(17:0), 55 pmol GlcCer(12:0), 14 pmol GM3 (18:0-D3), 359 pmol TG(47:1), 111 pmol CE(17:1), 64 pmol DG(31:1), 103 pmol MG(17:1), 724 pmol Chol(d6), 45 pmol Car(15:0)) were added and the sample sonicated for 2 min followed by centrifugation at 20000 g for 2 min. The supernatant was collected into a new tube and 200 µl chloroform and 800 µl 1% AcOH in water were added, the sample briefly shaken and centrifuged for 2 min at 20000 g. The upper aqueous phase was removed and the entire lower phase transferred into a new tube and evaporated in the speed vac (45°C, 10 min). Spray buffer (500 µl of 8/5/1 2-propanol/MeOH/water, 10 mM ammonium acetate) was added, the sample sonicated for 5 min and infused at 10 µl/min into a Thermo Q Exactive Plus spectrometer equipped with the HESI II ion source for shotgun lipidomics. MS1 spectra (resolution 280000) were recorded in 100 m/z windows from 250 – 1200 m/z (pos.) and 200 – 1700 m/z (neg.) followed by recording MS/MS spectra (res. 70000) by data independent acquisition in 1 m/z windows from 200 – 1200 (pos.) and 200 – 1700 (neg.) m/z. Raw files were converted to .mzml files and imported into and analysed by LipidXplorer software using custom mfql files to identify sample lipids and internal standards. For further data processing, absolute amounts were calculated using the internal standard intensities followed by calculation of mol% of the identified lipids.

### Tandem affinity purification of tagged Creld2

HEK293 cells were electroporated with plasmid DNA pcDNA3.1/Zeo(+) (Addgene) containing only the Strep-Flag (SF)-tag (mock), N-terminally tagged Creld2 (FS-C2) where the tag was integrated after the signal peptide or C-terminally tagged Creld2 (C2-SF). Gene blocks were used to clone the constructs (see Table S6). Primer used for cloning:

NheI-Kozak-mC2-SF-tag fw 5’-CTAGCTAGCGCCACCATGCACCTGCTGCTTGCA-3’,

5’-Creld2-KpnI TATGGTACCTCACAAATCCTCACGGGAGGG-3’,

KpnI-Flag rev 5’-TAGGGTACCTCACTTGTCGTCGTC-3’,

NheI-SF-tag fw 5’-CTAGCTAGCATGGGTGGAGGTTCTGGA-3’,

KpnI-SF-tag rev 5’-GGAGCTCTGGATGGTACCTCACTTGTC-3’.

Plasmids comprised a zeocin resistance cassette for selection. Cells transiently expressing C2-SF were harvested 72 h post electroporation for tandem-affinity-purification (TAP). For the generation of stably expressing cell lines plasmid DNA was linearized prior to electroporation and selected for stable expression by Zeocin (200 µg/ml) resistance for 4 weeks with validation of construct expression via immunoblotting against Creld2 and Flag (data not shown). For TAP cells were incubated on ice in freshly prepared, ice cold TAP-lysis buffer (1x Complete protease inhibitor (Roche), 1x Phosphate inhibitor cocktail (PIC) I (Sigma-Aldrich), 1x PIC II (Sigma-Aldrich), 0.5% NP40 in 1x TBS) for 20 min, centrifuged for 10 min at 10000 g at 4°C and supernatant was recovered. Subsequently, lysates were incubated with Strep-tactin Sepharose resin (IBA) for 2 h at 4°C on a rotation wheel. After incubation lysates were transferred onto Micro Bio-Spin columns (0.8 ml, BioRad), washed 3 times with TAP-wash buffer (1x PIC I, 1x PIC II, 0.1% NP40 in 1x TBS) and incubated with TAP-elution buffer (50 µM Desthiobiotin (IBA) in 1x TBS) at 4°C for 10 min prior to elution by centrifugation at 100 g for 10 sec. Eluates were incubated with anti-Flag M2 resin (Sigma-Aldrich) for 2 h at 4°C on a rotation wheel, washed once with TAP-wash buffer, twice with TBS and eluted by incubating samples with Flag elution buffer (200 µg/ml Flag peptide (Sigma-Aldrich) in 1x TBS) for 10 min following centrifugation for 10 sec at 2000 g. TAPs were performed in triplicates from cells harvested on different days.

### Processing of TAP proteins for mass spectrometry and analysis

Eluates were reduced with 10 mM Dithiothreitol (DTT) for 30 min at 56 °C followed by alkylation with 55 mM Chloroacetamide (CAA) for 30 min at RT in the dark. Subsequently, samples were boiled in laemmli buffer (1 x final concentration 2% SDS, 2 µM DTT, 5% (v/v) Glycerol, 50 µM Tris-HCL (pH 6.8, 0.01% (w/v) Bromophenol blue) at 95 °C for 10 min and run for approx. 1 cm in a 10% SDS-PAGE gel to separate purified proteins from the Flag peptide used for elution. Gels were fixed in fixation buffer (45 % H^2^O, 45 % MeOH, 10 % Acetic acid) for 1 h at RT and stained (fixation buffer, 0.05 % Coomassie-G250, filtered) for 1 h at RT followed by a first destaining in fixation buffer for 1 h prior to destaining O/N. Protein lanes were separated, cut into small pieces, washed twice with MS-wash buffer (50 mM Ammonium bicarbonate (ABC), 50 % Acetonitrile (ACN)) for 20 min, dehydrated with ACN for 10 min and dried in a speed vac for 20 min. Afterwards, samples were washed with 50 mM ABC for 15 min at RT and dehydration was repeated with speed vacuuming for 30 min. For in-gel-digest of proteins, gel pieces were soaked in digest solution (0,009 µg/µl Trypsin, 0,001 µg/µl LysC, 45 mM ABC) for 30 min at 4 °C, covered with 50 mM ABC and digested O/N at 37 °C. Supernatants were recovered, gel pieces incubated in extraction buffer (30 % ACN, 3 % Trifluoroacetic acid (TFA)) for 20 min at RT, supernatants were recovered and proteins were extracted for another time from gel pieces by incubation in 100 % ACN for 20 min at RT with recovery of supernatants. The recovered supernatants of individual samples were pooled, organic solvents were evaporated in a speed vac, samples were acidified with Formic acid (1 % final concentration), stage tipped and subjected to mass spectrometry.

### TAP mass spectrometry

Raw mass spectrometry data files were analysed using MaxQuant software (version 1.6.0.16). Further analysis of quantitated proteins was done with Perseus software (version 1.5.5.3). Data was cleaned by filtering out of reverse proteins, proteins only identified by site and contaminants, and filtered for valid proteins label-free quantification (LFQ) values identified in at least 2 technical replicates in minimum 1 condition. Further analysis of data was done using R software. Missing LFQ values were imputed using the DEP package (version 1.6.1) by firstly imputing of missing not at random (MNAR) values with 0, followed by a screen for falsely categorized MNAR values and re-imputation of missing at random (MAR) values using the k-nearest neighbors (knn) algorithm. For statistical analysis of differentially enriched (DE) proteins two-tailed t-test with unequal variance was applied to conditions containing Creld2 protein versus the respective mock control (acute or stable). Proteins with fold change > 0 (Creld2 containing conditions versus mock control) and p-value < 0.1 were considered differentially enriched.

### Patients and ethics

The study protocol conformed to the revised Declaration of Helsinki (Edinburgh, 2000) and was approved by the local Institutional Review Board (Ethik-Kommission am Universitätsklinikum Essen; file number 09-4252). All patients provided written informed consent for participation in the study before recruitment. Liver tissue and serum were collected from morbidly obese patients undergoing bariatric surgery. All enrolled patients underwent physical and ultrasound examinations, and a complete set of laboratory studies. Subjects reporting excessive alcohol consumption (>20 g/day for men or >10 g/day for women) and those with other known causes of secondary fatty liver disease (e.g., viral hepatitis, metabolic liver disease, toxic liver disease) were excluded from the study. Histological evaluation of NAFLD and classification as NAFL or NASH were performed by two experienced pathologists according to the method of Bedossa et al. (SAF score [30]).

## CRELD2 ELISA

Serum CRELD2 levels were measured using a human CRELD2 Elisa kit (Ray Biotech) according to the manufacturers’ instructions. **Immunoblotting**

### Human tissue samples

Liver tissue was directly lysed for 30 min on ice with RIPA Lysis and Extraction Buffer (Thermo Scientific) containing protease inhibitor cocktail and phosphostop (Roche). After centrifugation at 13 000 g for 15 min at 4 °C, protein concentration in the supernatant was measured using Bradford’s reagent (Bio-Rad).

### Mouse tissue samples

Tissue was snap frozen in liquid N^2^ and stored at −80 °C until usage. Tissue was homogenized in RIPA buffer containing PIC I and PIC II (Sigma-Aldrich) and Complete protease inhibitor (Roche) using a Precellys homogenizer and lysed for 30 min on ice. 10-20 µg of total protein were separated by SDS-PAGE and transferred onto a PVDF membrane (Immobilon-P transfer membranes 0.45 µm, Merck) via semi-dry blotting (Trans-Blot Turbo Transfer System, Bio-Rad). If not otherwise mentioned, membranes were incubated with antibodies diluted in TBST (0.1% Tween20) over night at 4°C on a rocker with the particular antibodies (see Table S7). Protein band intensities were acquired using ImageJ2. Proteins of one membrane were normalized to Actin and set relative to a wildtype control located on the membrane.

#### Statistical analysis and reproducibility

Data are shown as mean with individual values per mouse represented as circles, unless stated otherwise. Statistical significance was analysed with R using unpaired two-tailed t-tests, one-way and two-way ANOVA as indicated in the figure legends. The n value represents biological replicates. Significance was considered at P < 0.05. Experiments were repeated to ensure reproducibility of the observations. No statistical methods were used to predetermine sample size.

## Data availability

Gene expression data are currently being deposited in the GEO database. Lipid mass spectrometry data are deposited in a Github repository https://github.com/maccabaeus/Creld2-lipid-mass-spectrometry.git). The data that support the findings of this study are available from the corresponding author upon reasonable request.

## Acknowledgements

We thank Prof. Michael Hoch for his long-standing support of this project. We thank Thomas D. Rutkowski for feedback on the manuscript. We thank Cornelia Cygon and Melanie Thielisch for technical support, and Dr. Joachim Degen for help with the generation of *Creld2*^*eGFP/eGFP*^ mice. We would like to thank Prof. Margarete Odenthal (Institute for Pathology, University Hospital Cologne), Prof. Hideo A. Baba und Martin Schlattjan (Institute for Pathology, University Hospital Essen), and Prof. Johannes Haybäck (Institute for Pathology, University Hospital Magdeburg) for histological preparation and assessments for the validation cohort. We also thank Prof. Niedergethmann (Department for General- and Visceral Surgery, Alfried Krupp Hospital, Essen, Germany) and Prof. Hasenberg (Helios Hospital Niederberg) for sample collection of the validation cohort. AC was funded by the DFG CA267/14-1. EM and JLS were funded by the DFG under Germany’s Excellence Strategy EXC2151-390873048. EM is supported by the Daimler and Benz Foundation. This work was funded by the Fritz Thyssen foundation (to EM, Az.10.18.2.029MN).

## Contributions

EM and PK conceived the project. PK, EM, NB, AF, LB, FB performed experiments. PK, EM, NB, AF analyzed data. EM, PK, and RB supervised experiments and data analysis. TU and JLS provided help with RNA-seq experiments and the script for CoCena^2^ analyses. CT helped with lipidomic analyses. JPS and AC assisted in interpretation of human results. EM and PK wrote the manuscript.

---

**Supplemental Figure 1:**
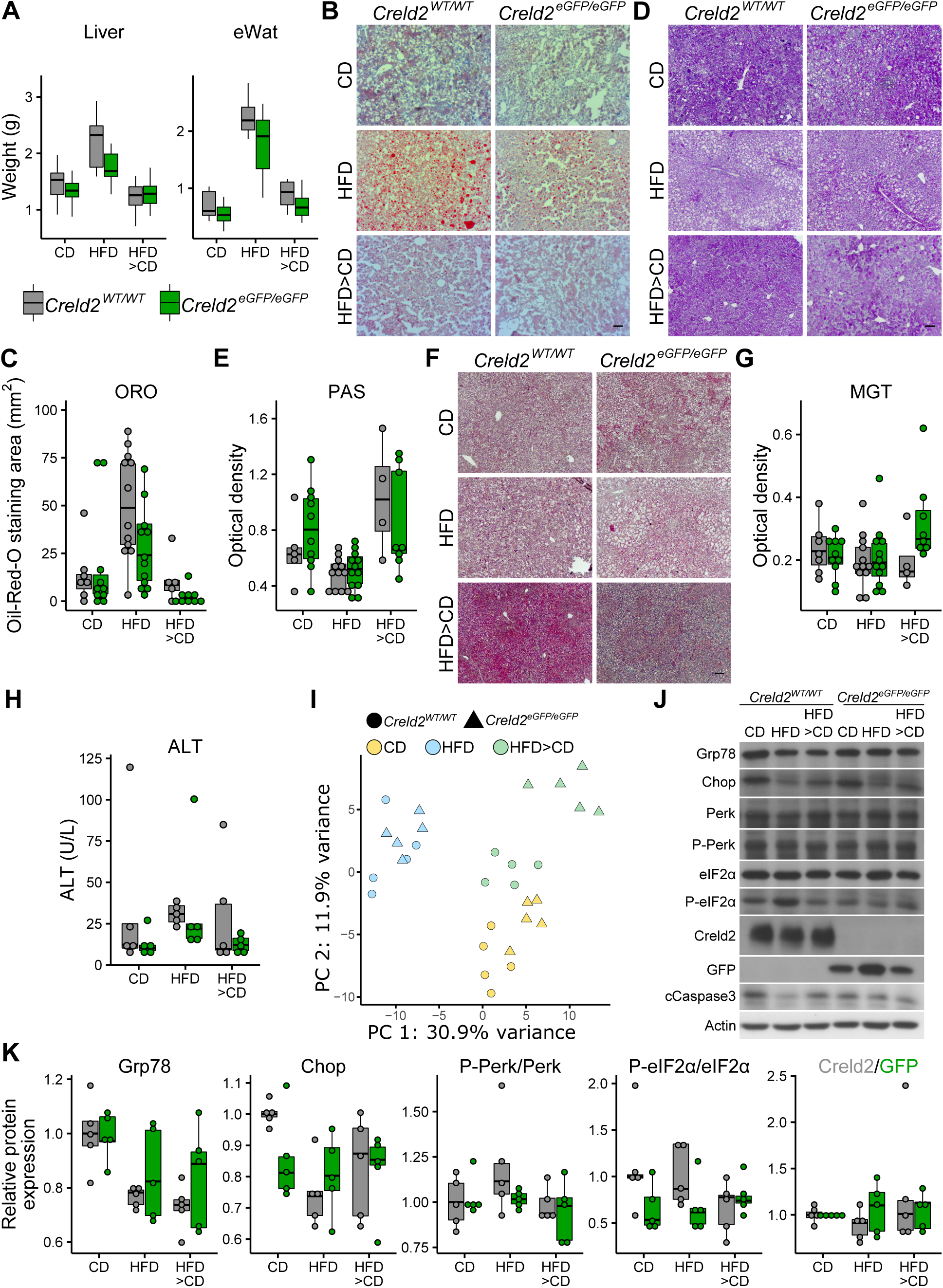
Diet-induced hepatosteatosis and analysis of liver damage. **(A)** Liver (n = 12-19) and eWat (epididymal white adipose tissue, n = 9-14) weights of mice after 12 weeks of diet. **(B-G)** Representative histological analyses of livers from mice on different diets by **(B)** ORO (oil-red-O), **(D)** PAS (Periodic Acid Schiff) and **(F)** MGT (Masson Goldner trichrome) staining and their corresponding quantification **(C, E, G). (H)** Determination of liver damage by serum alanine aminotransferase (ALT) concentration. Circles represent individual mice (C, E, G, H). Scale bars represent 100 µm (B, D, F). **(I)** Principle component analysis of liver RNA-seq data shown in Figure 2. **(J)** Representative immunoblots of liver protein lysates on different diets. Refers to main Figure 2H for cleaved Caspase 3. **(K)** Western blot quantification of UPR components (n = 5) from (J). For Perk and eIF2α, calculated using the formularatios of phosphorylated (P-Perk and P-eIF2α) and unphosphorylated protein abundance are displayed. Circles represent individual mice.

**Supplemental Figure 2:**
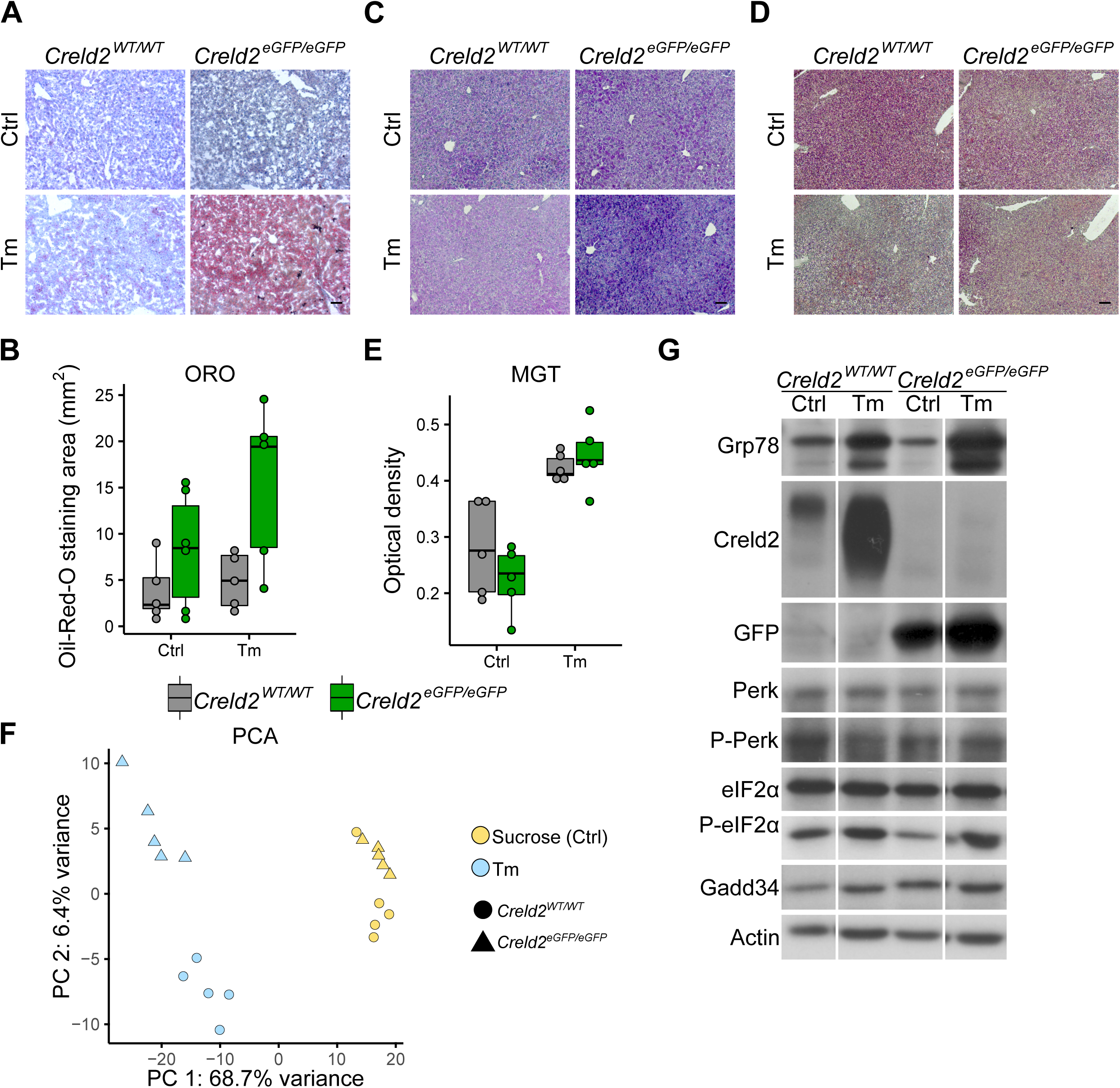
Pharmacologically induced hepatosteatosis and analysis of liver damage. **(A)** ORO (oil-red-O) staining of livers from ER stress induced mice (Ctrl: Sucrose control; Tm: Tunicamycin) and **(B)** ORO staining quantification. **(C)** PAS (Periodic Acid Schiff) staining. For PAS signal intensity quantification see Figure 3B. **(D, E)** MGT (Masson Goldner trichrome) staining and quantification. Circles represent individual mice (B, E). Scale bars represent 100 µm. **(F)** Principle component analysis of liver RNA-seq data shown in Figure 3. **(G)** Representative immunoblots of liver protein lysates injected with sucrose control (Ctrl) or Tm (tunicamycin). Refers to main Figure 3F.

